# Microbe-induced plant drought tolerance by ABA-mediated root morphogenesis and epigenetic reprogramming of gene expression

**DOI:** 10.1101/2023.01.03.522604

**Authors:** Khairiah M. Alwutayd, Anamika A. Rawat, Arsheed H. Sheikh, Marilia Almeida-Trapp, Alaguraj Veluchamy, Rewaa Jalal, Michael Karampelias, Katja Froehlich, Waad Alzaed, Naheed Tabassum, Thayssa Rabelo Schley, Anton R. Schaeffner, Ihsanullah Daur, Maged M. Saad, Heribert Hirt

## Abstract

The use of beneficial microbes to mitigate drought stress tolerance of plants is of great potential albeit little understood. We show here that a root endophytic desert bacterium, *Pseudomonas argentinensis* sp. SA190, enhances drought stress tolerance in Arabidopsis. Transcriptome and genetic analysis demonstrate that SA190-induced root morphogenesis and gene expression is mediated via the plant abscisic acid (ABA) pathway. Moreover, we demonstrate that SA190 primes the promoters of target genes in an epigenetic manner which is ABA-dependent. Application of the SA190 priming technology on crops is demonstrated for alfalfa in field trials, showing enhanced performance under desert agriculture conditions. In summary, a single beneficial root bacterial strain can help to perform agriculture under drought and water limiting conditions.

**Synopsis:** 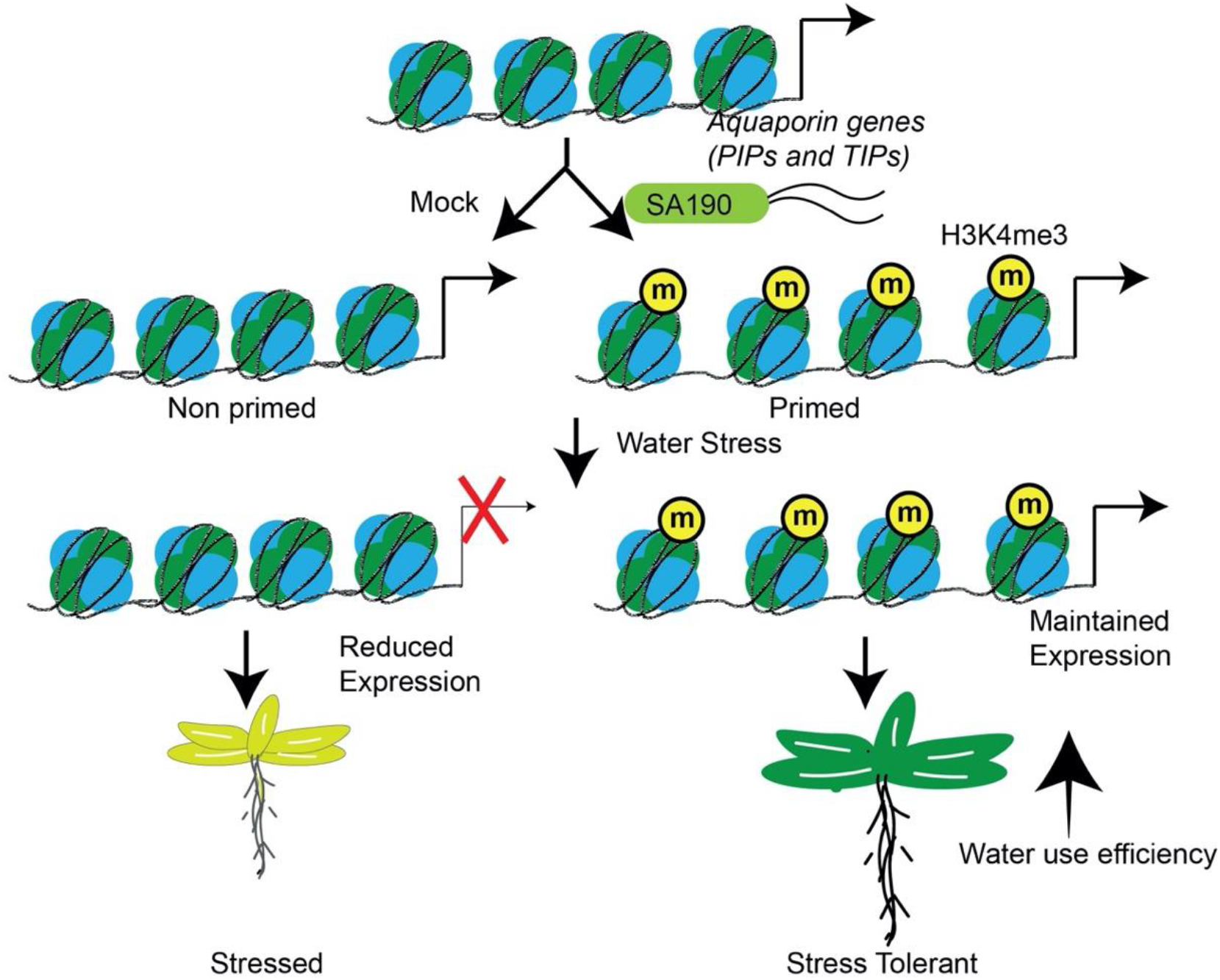

- Beneficial root endophyte *Pseudomonas argentinensis* sp. SA190 confers drought tolerance in plants
- SA190 modulates the expression of genes under drought stress in an ABA-dependent manner
- SA190 primes genes via H3K4me3 histone mark enrichment
- SA190 alters host plant physiology by improving the plant water status
- SA190 enhances crop performance in open field conditions with limited irrigation

## Introduction

Climate change increases the frequency, severity, and duration of drought in many parts of the world, making crop resistance to drought stress a major goal of agricultural biotechnology. The percentage of the planet affected by drought has doubled in the last 40 years which affected humanity more than any other natural hazard. Drought stress adversely affects the plant growth which results in reduced crop yield and ultimately to critical food shortages or famines (Seleiman et al, 2021). Although enormous progress has been made to understand and identify important molecular players of drought stress in model plants, success in knowledge transfer to economical crops is still limited (Cominelli et al, 2013). This is partially due to the fact that drought stress is a complex trait causing dramatic changes in many physiological plant parameters (Zhu, 2002). After all, the different stress conditions that coexist in open field agriculture go beyond simple arithmetic of single stress tolerance traits (Suzuki et al, 2014).

Drought stress drastically changes plant morphology, biochemistry and physiology leading to sever impact on plant growth and yield. The first physiological response of plants to drought is the closure of stomata to avoid water loss via transpiration (Pirasteh-Anosheh et al, 2016). The decrease in transpiration rate leads to a decrease in relative water content (RWC) of plants (Giday et al, 2014). Stomatal closure also decreases plant photosynthetic efficiency by decreasing intracellular CO2 concentrations (Lisar et al, 2016).

Abscisic acid (ABA) is one of the key phytohormones that regulates the plant response to drought stress. ABA acts through a conserved signal transduction pathway, comprised of a PYRABACTIN RESISTANCE 1-Like (PYL)-PROTEIN PHOSPHATASE 2C (PP2C) and SNF1-RELATED PROTEIN KINASE 2 (SnRK2) module. ABA binding to PYL protein triggers a conformational change in receptors which allows it to bind and inhibit the PP2C that normally represses ABA signaling. This PYL–ABA–PP2C complex releases SnRK2 from the otherwise inhibitory complex with PP2C, initiating phosphorylation of transcription factors which regulate gene expression involved in ABA output responses (Chen et al, 2020; Cutler et al, 2010; Fujii et al, 2009). ABA also regulates the early production of reactive oxygen species (ROS) after drought perception (Cruz de Carvalho, 2008). ROS serves as a stress signal to activate downstream processes, but accumulation of ROS can lead to cell damage and growth penalty.

Water movement across the cellular membranes is largely regulated by a family of water channel proteins called aquaporins. In Arabidopsis there exist 35 aquaporin genes (Johanson et al, 2001) which can be broadly divided into plasma membrane intrinsic proteins (PIPs) and tonoplast intrinsic protein (TIPs) genes. Some of the Arabidopsis PIPs and TIPs proved to be active water channels in *Xenopus* oocytes (Quigley et al, 2002). The biological significance of aquaporins in plants is their ability to modulate transmembrane water transport in situations where adjustment of water flow is physiologically critical (Li et al, 2014). PIPs play an important role in controlling the transcellular water transport and are subdivided into the two subfamilies PIP1 and PIP2 (Maurel et al, 2015). The overexpression of PIP isoforms increased the root osmotic hydraulic conductivity, transpiration and shoot to root ratio while the downregulation leads to drought stress susceptibility (Pawłowicz & Masajada, 2019). However, when expressed in a heterologous systems, the overexpression of aquaporins can lead to a negative effect on stress resistance, due to the fact that the native stress response machinery may recognize them as a foreign proteins (Li et al., 2014).

The transcriptional responsiveness of drought stress regulated genes is correlated with changes in histone modifications (Kim et al, 2012; Kim et al, 2008; To & Kim, 2014). Two of the important histone modifications, H3K4me3 and H3K9ac, are enriched on drought stress-regulated genes like *RD20* and *RD29A*. The levels of these modifications change from mild to severe drought stress, suggesting that epigenetic responsiveness depend on the intensity of the drought stress (Kim et al., 2012; Kim et al., 2008).

Beneficial plant microbes have been used to overcome abiotic stress challenges of plants (Busby et al, 2017; de Zelicourt et al, 2013). Indeed, plants and their rhizosphere host diverse microbial communities, selected from bulk soil (Bulgarelli et al, 2012), and the beneficial bacteria, defined as Plant Growth-Promoting Bacteria (PGPB), can establish symbiotic associations to promote plant growth under optimal conditions or in response to biotic and abiotic stresses (de Zelicourt et al, 2018; Finkel et al, 2017; Saad et al, 2020; Synek et al, 2021). A number of PGPBs have been reported to mitigate drought stress in a variety of plant species implicating a number of processes, such as modification of phytohormonal levels, production of heat-shock proteins and dehydrins, production of osmolytes, exopolysaccharides, volatiles, detoxification of reactive oxygen species (ROS) and the accumulation of compatible solutes like sugars, amino acids and polyamines (Sati et al, 2022). However, a detailed molecular and genetic analysis of the underlying molecular mechanisms has not been done so far.

The ideal natural reservoir for the isolation of beneficial bacteria which help in drought stress tolerance is the desert. To this end, a number of rhizo- and endosphere bacterial strains from desert plants were isolated and abiotic stress screens were conducted on Arabidopsis under controlled lab conditions (Bokhari et al, 2020; de Zelicourt et al., 2018; Lafi et al, 2016). Here, we report on *Pseudomonas argentinensis* sp. SA190, an endophytic bacterium isolated from root nodules of the indigenous desert plant *Indigofera argentea* (Lafi et al., 2016), that massively increases drought tolerance in plants. Physiological and genetic analysis showed that SA190 maintains growth and photosynthesis by enhancing plant water use efficiency. Transcriptome analysis uncovered that SA190 induces the expression of multiple aquaporin genes in roots of Arabidopsis. Inhibition of aquaporin function abrogates SA190-induced plant drought tolerance. SA190 epigenetically regulates gene expression by enhanced H3K4me3 deposition. The agronomic potential of SA190 is shown in field trials with alfalfa (*Medicago sativa*) under desert agriculture. SA190 colonized plants surpassed the productivity of alfalfa under limited irrigation, demonstrating that SA190 is a powerful agent to reduce water consumption in agriculture.

## Results

### *Pseudomonas argentinensis sp. SA190* enhances drought tolerance of Arabidopsis

To identify desert PGPBs with the potential to enhance drought stress tolerance in plants, we set up a screening system using the model genetic plant *Arabidopsis thaliana* ecotype Col-0 by assessing the capacity of various dessert root endophytic microbial strains to maintain plant growth on media infiltrated with 25% PEG as a proxy for drought stress. We identified *Pseudomonas argentinensis sp*. SA190 as one of the top candidates in the screen. To quantify the effect of SA190 under drought mimicking conditions, plants were germinated for 5 days on ½ MS agar medium containing 10^8^ cfu of *Pseudomonas argentinensis sp*. SA190 before transfer to fresh ½ MS plates infiltrated with 0 or 25% PEG (Fig. 1A). After 16 days, plant morphology, total fresh and dry weight, root length and lateral root density were determined (Fig. 1). Our results show that SA190 did not significantly influence the morphology or development of Arabidopsis under non-stress conditions (Fig. 1B). Upon 25% PEG treatment, the growth of non-colonized plants was severely inhibited (Fig.1B) resulting in a reduction of fresh and dry weight by more than 90% (Fig. 1C-D). In contrast, Arabidopsis colonized with SA190 maintained growth, resulting in a 6-fold enhancement of fresh and dry weight on 25% PEG (Figs. 1C-D, EV1).

**Figure 1.**
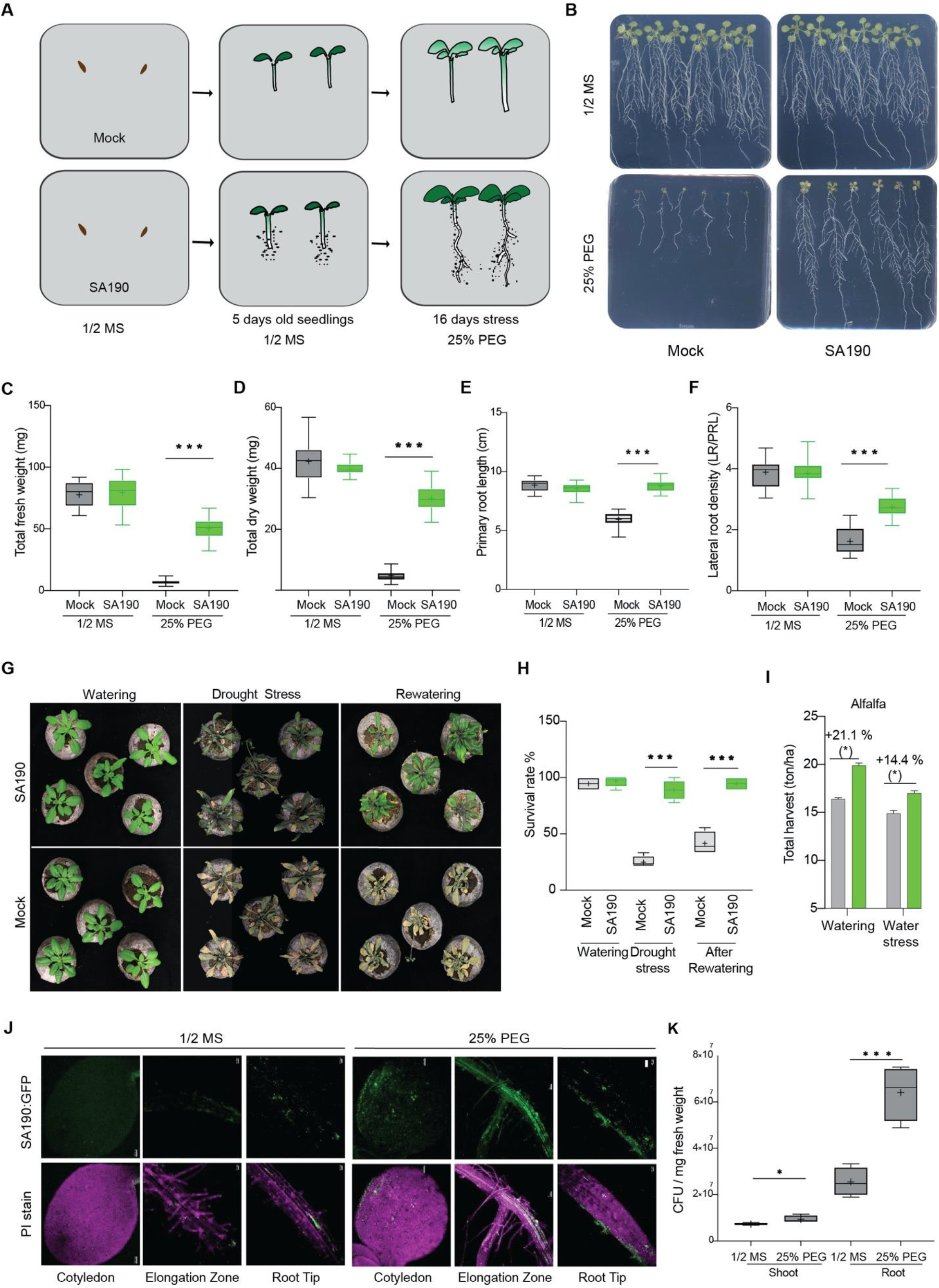
*Pseudomonas argentinensis* sp. SA190 enhances drought tolerance in Arabidopsis and crops. **A.** Graphical representation of work flow for plant screening assay under 25% PEG. **B.** Growth of non-colonized and SA190-colonized 21-day-old Arabidopsis plants grown under drought stress (½ MS + 25% PEG). **C.** Total fresh weight, **D.** Total dry weight, **E.** Primary root length and **F.** Lateral roots density of 21-day-old, mock- and SA190-colonized plants upon growth for 16 days on ½ MS or ½ MS + 25% PEG. **G.** Growth and **H.** Survival percentage of mock- and SA190-colonized plants in jiffy pots grown upon watering for 2 weeks, no watering for 3 weeks and after one week of re-watering. All plots represent the mean of 3 biological replicates (n= 36). Error bars represent SE. Asterisks indicate the statistical differences based on the Student’s t-test (* P < 0.05; ** P < 0.01; *** P < 0.001). **I.** Alfalfa yield in sandy soil open field agriculture with normal or reduced irrigation (30% reduction). The differences in yield between different treatments are indicated in percent (%) and significance at P<0.05 is indicated by asterisks (*). **J.** Confocal microscopy showing the colonization of SA190:GFP on five days-old seedlings grown on ½ MS and 25% PEG. Green signal corresponds to GFP-tagged SA190 while in magenta are plant cell wall stained with propidium iodide (PI). Bar indicates 100 nm. **K.** SA190 colonization levels in Arabidopsis under normal and drought conditions as quantified by colony forming units of bacteria per mg of plant fresh weight.

### *Pseudomonas argentinensis sp. SA190* enhances drought tolerance of Arabidopsis and crops in desert agriculture

To see the effect of SA190 on the growth of adult plants under drought conditions, 5 day-old non-colonized or SA190-colonized Arabidopsis were transferred to pots for two weeks. Thereafter, watering was stopped, and plants were subjected to drought stress for three weeks. While non-colonized (Mock) plants showed a drop in survival by more than 70%, SA190-colonized plants were affected by less than 10% (Fig. 1G-H). One week after re-watering, less than 50% of non-colonized plants resumed growth in comparison to a 100% of SA190-colonized plants (Fig. 1G-H). These results indicate that SA190 significantly enhances drought stress tolerance of Arabidopsis plants.

To evaluate the agronomic potential of SA190 on crops, we performed open field trials with alfalfa which is used as an important animal feed in different regions of the world. Alfalfa seeds were coated with SA190 and tested in parallel with mock-coated seeds in open field desert agriculture. A randomized complete block design with a split-split plot arrangement with different replicates was used with a water irrigation protocol to induce mild drought stress. SA190-inoculated alfalfa plants exhibited a significant yield gain under water stress conditions (Fig. 1I). These results show that SA190 can efficiently enhance the performance of crops in open field conditions. SA190 enhanced productivity of alfalfa also under normal irrigation (Fig. 1I), which can be explained by the fact that the desert agriculture production conditions are far from optimal for crop growth and yield.

### SA190-induced root morphogenesis and enhanced root colonization of Arabidopsis upon drought stress

Since SA190 was isolated from roots of the desert plant *Indigofera argentea*, we characterized the interaction between SA190 and Arabidopsis in more detail. To this end, SA190 was stably transformed with a GFP expressing cassette (SA190:GFP). Confocal microscopy revealed that SA190:GFP mainly colonized roots (Fig. 1J), preferentially epidermal cells of the root elongation zone (Fig. 1J). Interestingly, the roots of plants grown on 25% PEG showed enhanced colonization when compared to plants grown only on ½ MS agar (Fig. 1J). These findings were confirmed by determining the SA190 colony forming units (CFU) of root and shoot tissues (Fig. 1K).

We next investigated the root morphology of SA190-colonized plants. After 16 days on normal or 25% PEG medium, root length and lateral root density were determined. Our results show that SA190 did not significantly influence the morphology or development of Arabidopsis roots under non-stress conditions (Fig. 1B, EV1B, EV1D). However, on 25% PEG medium and in contrast to non-colonized plants, SA190-colonized plants showed a strongly developed root system (Fig. 1B), with a massive increase in root fresh and dry weight (EV1B, EV1D) mostly due to enhanced growth of the primary root and development of the lateral root system (Fig. 1E-F).

### SA190 reprograms the transcriptional drought stress response of Arabidopsis

To understand the molecular mechanism of SA190 induced drought tolerance, we next performed RNA-seq analysis of 21-day old non- or SA190-colonized roots upon normal and drought stress for 16 days. The root transcriptome data were organized by hierarchical clustering into groups of differentially expressed genes (DEGs) according to their expression patterns in non-colonized (Mock) and SA190 colonized plants (SA190) under normal conditions, as well as non-colonized and SA190-colonized plants under drought stress conditions, denoted as PEG and PEG+SA190, respectively (Fig. 2A). We then assessed the gene ontology enrichment of the up- and down-regulated root DEGs. When comparing SA190- to mock-colonzied plants, we only observed 30 up- and 145 down-regulated DEGs under non-stress conditions. In this relatively small gene set, we nonetheless found significant GO term enrichment of genes related to cell wall organization (eg. expansins and pectin lyases) and response to water deprivation.

**Figure 2.**
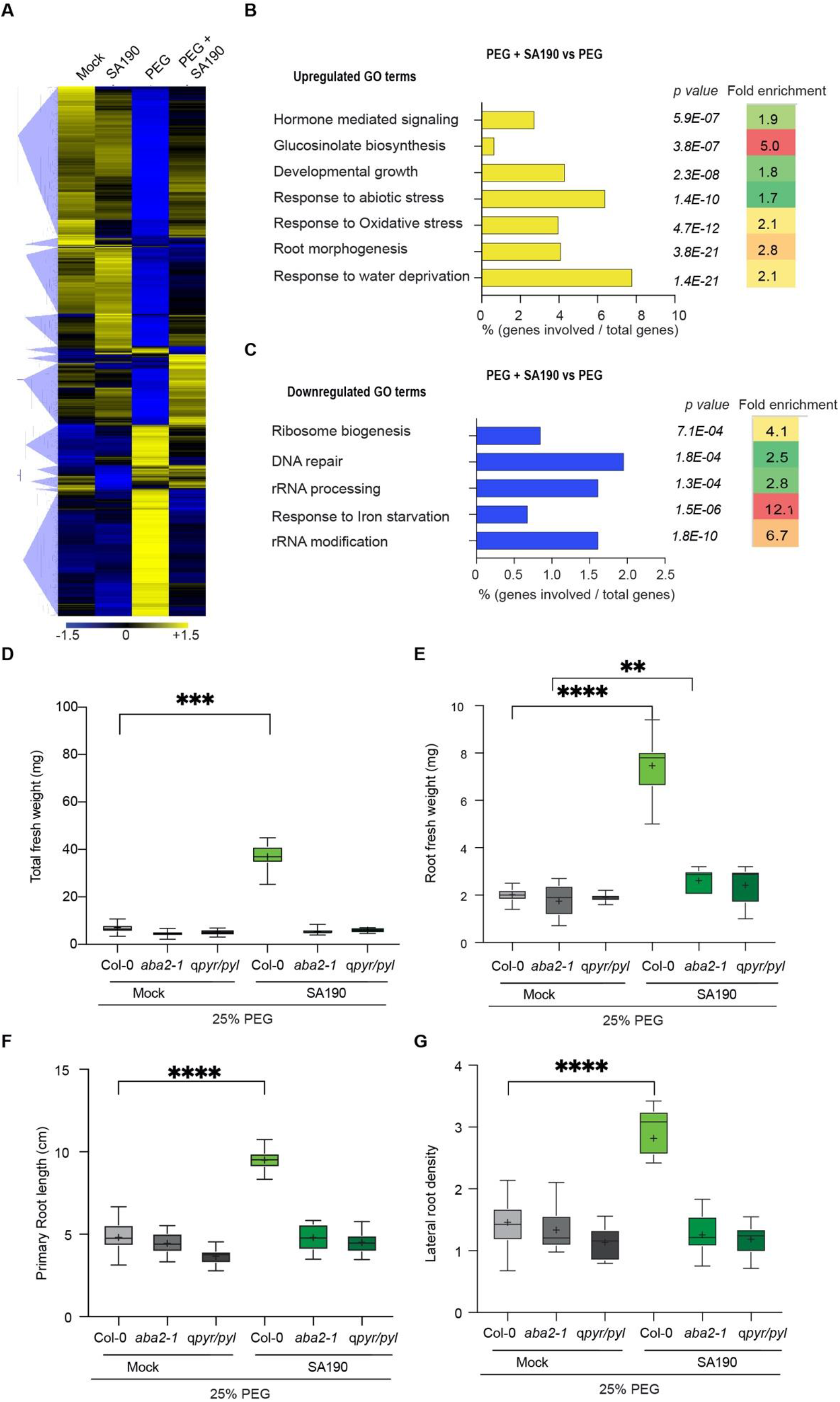
Transcriptome analysis and role of ABA signaling in SA190-induced drought tolerance. **A.** Heatmap showing hierarchical clustering of up- and down-regulated DEGs profiles in response to PEG (25%), SA190 or both treatments based on root RNA-Seq analysis. Quantified read counts are normalized for genes. Colors indicate normalized expression levels. q < 0.05 and a fold change cut-off was set to log_2_>1 for up- and log_2_>-1 for down-regulated genes. **B-C.** GO term analysis of differentially expressed genes of SA190-colonized (PEG+190) compared with non-colonized (PEG) plants under drought stress conditions. **D-E.** Total fresh weight and root fresh weight of 21 days old non-colonized- (Mock) and SA190-colonized (SA190) Col-0, *aba2-1* and *qpyr/pyl* plants grown drought conditions (½ MS + 25% PEG). **F-G.** Primary root length and Lateral root density of 21-day-old, mock- and SA190-colonized Col-0, *aba2-1* and *qpyr/pyl* plants upon growth for 16 days on ½ MS + 25% PEG.

Under drought stress conditions, massive changes in DEGs were seen between SA190-colonized and non-colonized roots revealing 2384 up and 1365 down-regulated genes. Whereas the up-regulated genes showed enrichment in responses to hormone signaling, glucosinolate biosynthesis, water deprivation and oxidative stress as well as root morphogenesis and developmental growth (Fig. 2B). The most significant GO terms in the set of down-regulated genes were related to ribosome, RNA metabolism and DNA repair (Fig. 2C). Interestingly, the hormone, abiotic stress and water deprivation-related GO terms were all related to ABA signaling, including the ABA receptor components PYL1, PYL3, the ABA protein phosphatases HAI1, and HAI2, as well as the key ABA responsive protein kinase SnRK2-3 and a large number of aquaporin genes.

### ABA pathway mutants are compromised in SA190-induced drought tolerance

Since ABA plays a key role in drought stress tolerance and our transcriptome analysis showed that SA190 colonization of plants affects ABA signaling under drought stress, we tested Arabidopsis mutants impaired in ABA biosynthesis (*aba2-1*) and signaling *pyr1 pyl4 pyl5 pyl8* (*qpyr/pyl*). Under non-stress conditions, we observed no significant differences in the fresh or dry weight of SA190-colonized and non-colonized plants between WT and both mutants (Fig. EV3A). Under drought stress, however, SA190-maintained plant growth was strongly compromised in both *aba2-1* and *qpyr/pyl* mutants (Fig. 2D), indicating a central role of the ABA pathway in the SA190-induced Arabidopsis drought stress tolerance.

### SA190-induced changes in root morphogenesis are dependent on ABA signaling

Since ABA plays an important role in pathogenic plant-microbe interactions, we tested whether ABA-deficient mutants might compromise the interaction of Arabidopsis with SA190 and therefore result in the loss of the beneficial effect on drought stress tolerance. However, similar levels of SA190 colonization were seen under non-stress (½ MS) and drought stress (½ MS+25%PEG) conditions (Fig. EV2), indicating that ABA does not influence SA190 colonization of Arabidopsis.

Since SA190 induces massive changes in Arabidopsis root structure, we next tested whether ABA pathway mutants affect SA190-induced modification of Arabidopsis root morphogenesis. Under non-stress conditions, we observed no significant differences in the fresh weight of SA190-colonized and non-colonized roots when comparing WT to either ABA biosynthesis (*aba2-1*) or ABA signaling *pyr1 pyl4 pyl5 pyl8* (*qpyr/pyl*) mutants (Fig. EV3B-D). Under drought stress, however, the SA190-induced changes in root structure, including fresh weight, primary root length and lateral root density were strongly compromised in both *aba2-1* and *qpyr/pyl* mutants (Fig. 2 E-G), indicating a central role of the ABA pathway in SA190-induced changes in root morphogenesis.

### SA190 modulates the expression of aquaporin genes under drought stress in an ABA-dependent manner

RNAseq analysis of root tissue revealed that under drought stress conditions, a large set of Arabidopsis genes including several aquaporins were differentially regulated by SA190 (Fig. 3A). Since aquaporins are critical elements in adjusting of water flow in physiologically critical situations (Li et al., 2014), many of which are regulated by ABA (Pawłowicz & Masajada, 2019), we analyzed these aquaporin genes as proxy for the set of SA190-regulated genes in more detail. qRT-PCR analysis of aquaporin transcript levels did not change under non-stress conditions, but showed significantly higher levels in roots of SA190-colonized compared to non-inoculated plants (Fig. 3B). Both *aba2-1* and *qpyr/pyl* mutants strongly compromised the SA190-enhanced expression of all tested aquaporin genes under drought stress conditions (Fig. 3C), indicating that the ABA pathway is essential in mediating the SA190-induced changes of aquaporin gene expression.

**Figure 3.**
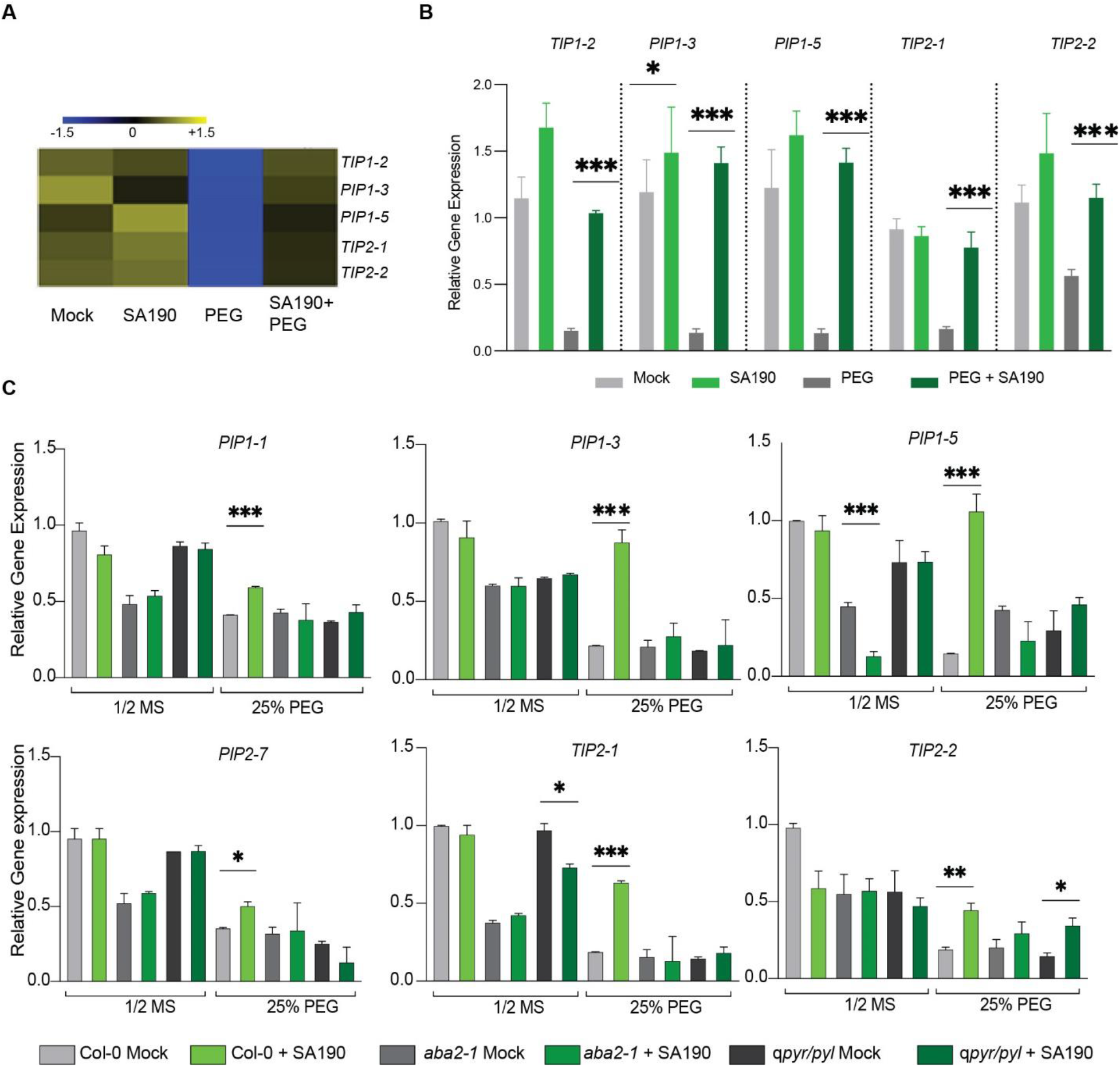
SA190-induced aquaporin gene expression is mediated in an ABA-dependent manner. **A.** Heat map of expression profile of differentially regulated *PIP* and *TIP* genes in root RNA-Seq analysis. **B.** Relative gene expression of aquaporin *PIP* and *TIP* genes by qRT-PCR analysis normalized to *tubulin* levels in 21-days old mock- and SA190-colonized roots of Arabidopsis grown for 16 days on ½ MS with or without 25% PEG. **C.** Relative gene expression of aquaporin *PIP* and *TIP* genes by qRT-PCR analysis in Col-0, *aba2-1* and *qpyr/pyl* mutants normalized to *tubulin* levels in 21-days old mock- and SA190-colonized roots of Arabidopsis grown for 16 days on ½ MS with or without 25% PEG. Values represent the means of three biological experiments. Error bars indicate SE. All plots represent the mean of 3 biological replicates (n= 36). Asterisks indicate a statistical difference based on the Student’s t-test (* P < 0.05; ** P < 0.01; *** P < 0.001).

### Epigenetic priming of SA190-targeted genes is mediated by the ABA pathway

Our transcriptome analysis of SA190-colonized plants showed that the expression of many genes is not constantly increased, but only shows differential changes under drought stress conditions. We therefore tested whether SA190 colonization might already be related to changes in the epigenetic state of the differentially regulated aquaporin genes under non-stress conditions. As H3K4me3 was found as a priming mark for genes in drought-trained plants (Ding et al, 2012a), we tested different regions of the set of SA190-regulated aquaporin genes for H3K4me3 enrichment. We observed a significant enrichment of the H3K4me3 mark in these aquaporin genes, especially near transcription start sites (Fig. 4). Since ABA was essential for mediating SA190-induced aquaporin gene expression and drought tolerance, we also analyzed the epigenetic status of these aquaporin genes in the ABA-deficient *aba2-1* mutant. In non-colonized (mock) plants, the *aba2-1* mutant showed similar levels of H3K4me3 as wild type plants. In SA190-colonized plants, however, *aba2-1* mutant plants were completely compromised in the enhancement of H3K4me3 levels in the aquaporin gene loci. We conclude that SA190 primes aquaporin genes in an ABA-dependent manner by enhancing H3K4me3 levels.

**Figure 4.**
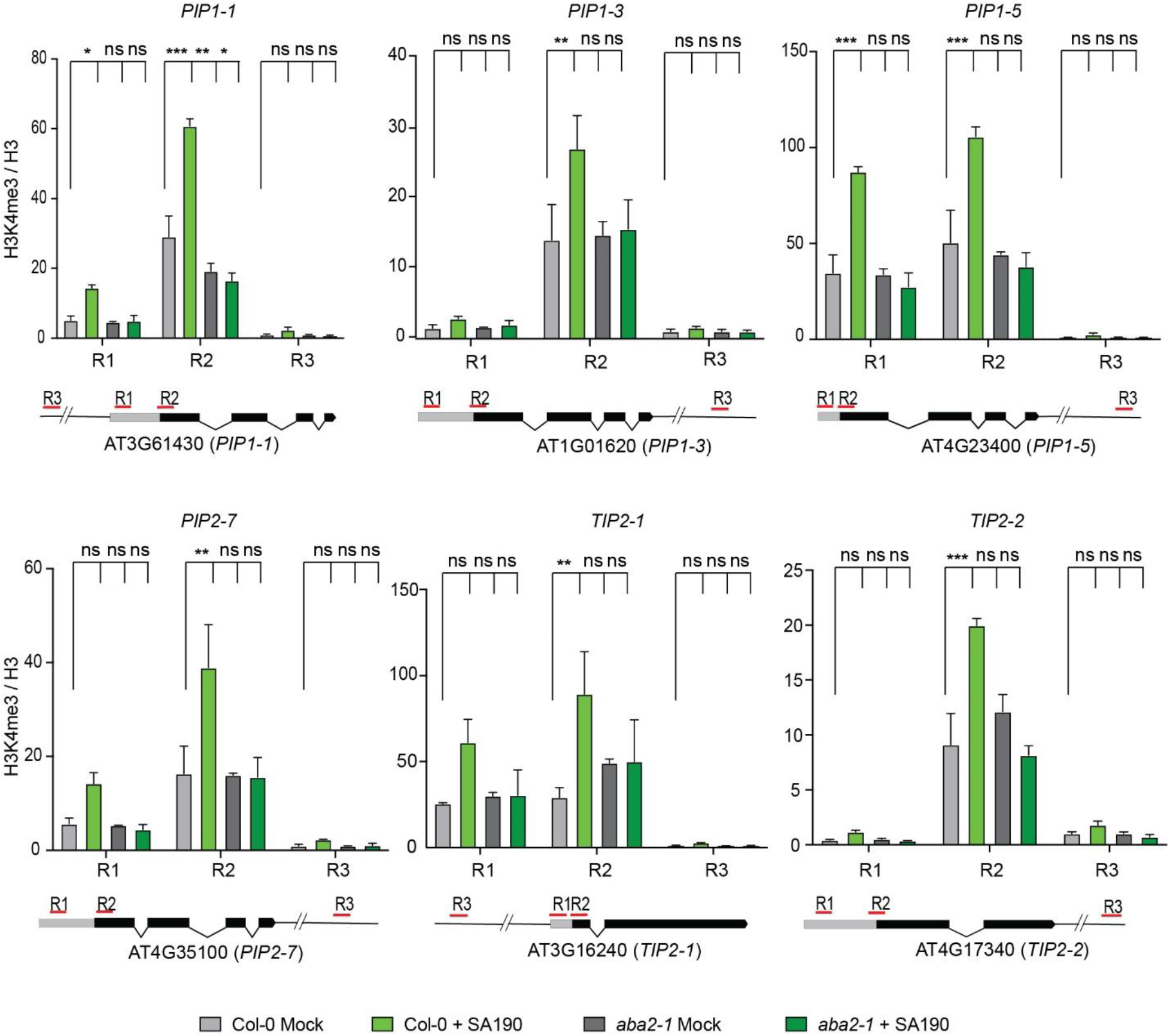
SA190 primes aquaporin genes via H3K4me3 enrichment. Relative enrichment of the H3K4me3 mark at the indicated gene loci of selected *PIP*s and *TIPs* of non-colonized (Mock) and SA190-colonized (SA190) plants in Col-0 and *aba2-1* mutant as determined by chromatin immunoprecipitation-qPCR (ChIP-qPCR). The regions targeted for amplification are labelled as R1, R2 and R3. R1 encompasses 5’-UTR, R2 is close to the transcription start site (TSS) while R3 is outside the gene body. Amplification values were normalized to input and H3 and region 3 (R3) of non-colonized (Mock) Col-0 plants. The plots represent the means of 2 biological replicates. Error bars represent SE. (ns) non-significant Asterisks indicate a statistical difference based on 2-way ANOVA (\**P* ≤ 0.05; \*\**P* ≤ 0.01; \*\*\**P* ≤ 0.001 for P value differences between the conditions when compared to Col-0 Mock using Dunnett’s multiple comparison test).

### Role of aquaporins in SA190-induced drought stress tolerance

Our analysis suggested that aquaporins might be key targets and mediators of SA190-induced drought stress tolerance. Aquaporins play a key role in transmembrane water transport in environmental stress responses (Li et al., 2014; Maurel et al., 2015) and their enhanced expression increased the root osmotic hydraulic conductivity, transpiration and shoot to root ratio while the downregulation lead to higher drought stress susceptibility (Pawłowicz & Masajada, 2019). To directly test the significance of the set of aquaporins identified in the SA190 transcriptome in drought stress, we used the quintuple *pip1;1 pip1;2 pip1;3 pip1,4 pip1;5* (*qpip*) and *tip2;1* Arabidopsis mutants in the absence and presence of SA190 (Fig. 5A-B). No significant changes of SA190-idnuced drought tolerance were observed when comparing growth of WT with *qpip* and *tip2;1* mutants under non-stress (Fig. 5A) or drought stress conditions (Fig. 5B). We also assessed whether the aquaporin mutants affect SA190 colonization. Although drought clearly induced SA190 colonization of Arabidopsis, this effect was similar for WT, *qpip* and *tip2;1* mutants (Fig. 5C).

**Figure 5.**
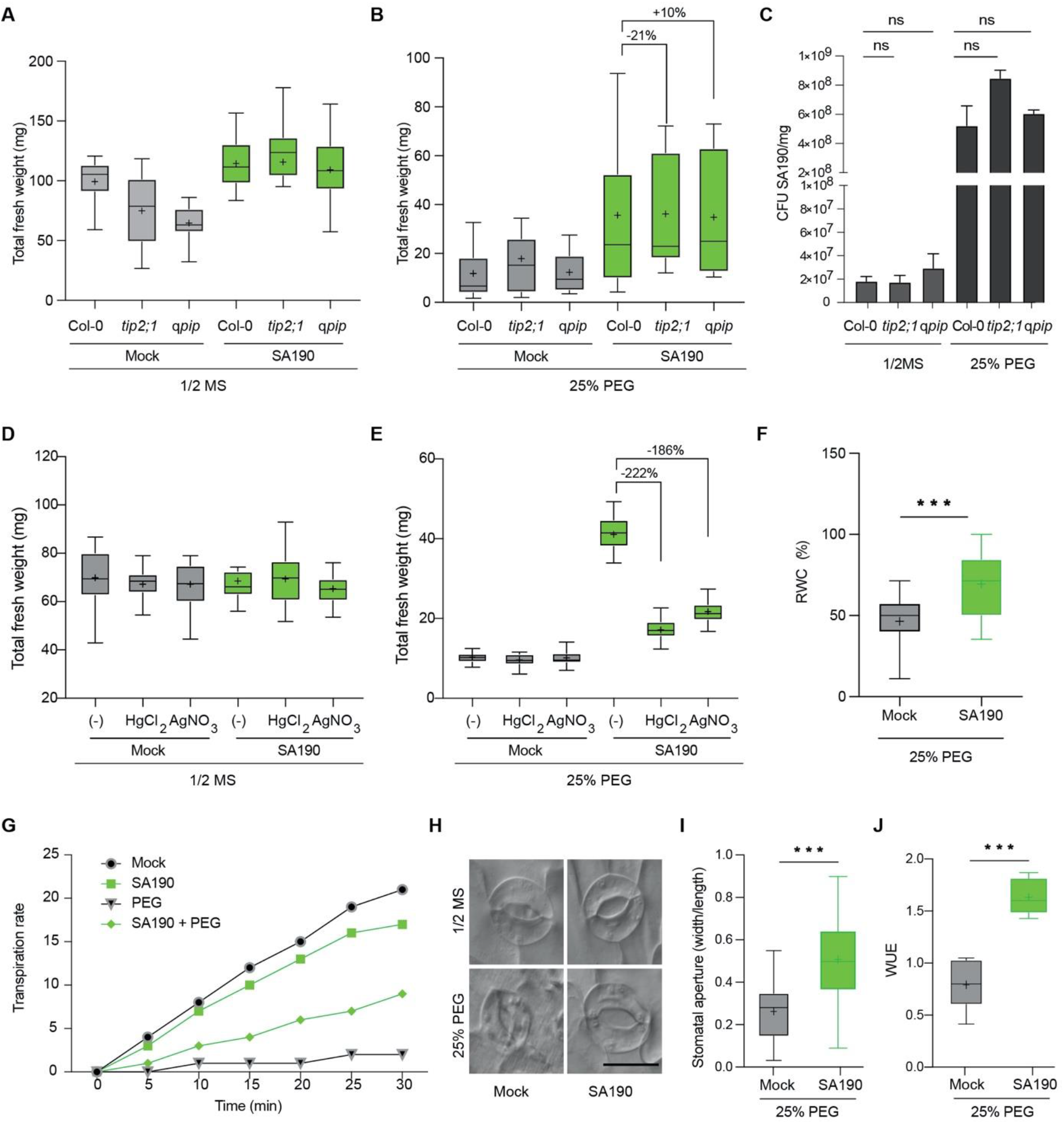
Role of SA190 colonization in aquaporin and plant drought stress physiology. Role of aquaporins in SA190-induced drought tolerance. **A-B.** Total fresh weight of 21 days old non-colonized-(Mock) and SA190-colonized (SA190) Col-0, *tip2;1* and *qpip* plants grown for 16 days under normal (½ MS) or drought conditions (½ MS + 25% PEG). **C.** Quantification of SA190 colonization efficiency in 21 days old - SA190-colonized (SA190) Col-0, *tip2;1* and *qpip* plants grown for 16 days under normal (½ MS) or drought conditions (½ MS + 25% PEG). **D-E.** Total fresh weight of mock- and SA190-colonized 21-day-old Arabidopsis plants grown for 16 days under normal on (½ MS) or drought (½ MS + 25% PEG) conditions supplemented with the aquaporin inhibitors HgCl_2_ (38.2 μM) or AgNO_3_ (5 μM). **F.** The percentage of leaf relative water content (RWC) of 21-days old mock- or SA190-colonized plants grown for 16 days under drought stress (1/2MS + 25% PEG) conditions. The RWC was calculated as the formula RWC (%) = [(FW-DW)/(TW-DW)]x100, where FW is fresh weight, DW is dry weight and TW is the weight of fully turgid leaves. **G.** Transpiration rates of mock- or SA190-colonized 21 days old plants after transfer of 5-day-old seedlings from ½ MS to ½ MS +/-25 % PEG. Transpiration rate was quantified as rate of water loss by calculating the changes in fresh weight for 30 min with reading at every 5 min interval. **H-I.** Comparison of stomatal aperture of SA190- and non-colonized 21-days old plants grown for 16 days under non-stress (½ MS) or drought stress (½ MS + 25% PEG) conditions. Scale bar = 20 μm. **J.** Water use efficiency (WUE) of 4-weeks old mock- or SA190-colonized plants under drought stress conditions. WUE was calculated as mg of dry weight produced per ml of water used. All plots represent the mean of 3 biological replicates. Error bars represent SE. Asterisks indicate a statistical difference based on the Student’s t-test (* P < 0.05; ** P < 0.01; *** P < 0.001).

The maintenance of the beneficial response of Arabidopsis to SA190 in the *pip* and *tip* mutants might also be due to the redundancy of the large aquaporin gene family in Arabidopsis. To overcome this problem, we also made use of two known aquaporin inhibitors, AgNO_3_ and HgCl_2_, which had no effect on Arabidopsis growth under normal conditions, irrespectively whether the plants were SA190- or non-colonized (Fig. 5D). However, under drought stress conditions, the inhibitors completely compromised the beneficial effect of SA190 on Arabidopsis (Fig. 5E).

The fact that one of the aquaporin inhibitors was AgNO_3_, which is also an inhibitor of ethylene signaling, made us question its role in the beneficial interaction of SA190 with Arabidopsis. We therefore tested a number of ethylene biosynthesis and signaling mutants for a possible role in the SA190-Arabidopsis interaction. None of the ethylene mutants affected the growth of SA190-colonized plants in comparison to non-colonized plants under non-stress conditions (Fig. EV4A). Compared to the massive loss of beneficial activity of SA190 in ABA mutants (Fig. 2D), SA190-induced drought stress tolerance in the ethylene mutants was only mildly affected under water limiting conditions (Fig. EV4B), suggesting that ethylene does not play a major role in mediating SA190 drought stress tolerance of Arabidopsis plants.

### SA190 colonization changes plant drought stress physiology

The importance of aquaporins in regulating transpiration and photosynthesis in drought stress and its importance in water use efficiency, growth and yield was recently highlighted (Moshelion et al, 2015). We therefore addressed whether similar effects could be observed in SA190-colonized plants when exposed to drought stress conditions.

Since drought stress leads to a decreased water content in plants, we measured the water content of leaves in SA190-colonized and non-colonized plants under non-stress and drought conditions. Under non-stress conditions, no significant differences in the relative water content (RWC) of leaves were observed when comparing SA190- and non-colonized plants (Fig. EV5A). Drought stress caused a massive reduction in the relative leaf water content in non-colonized plants (Fig. 5F), while SA190-colonized plants maintained leaf relative water content to the same level as seen in plants grown under non-stress conditions (Fig. EV5A).

Plants commonly react to drought stress by reducing transpiration to save water. However, reduced transpiration also means reduced photosynthesis and growth. Under non stress conditions, we observed a slightly lower transpiration rate in SA190-colonized plants when compared to non-colonized plants (Fig. 5G). In contrast, whereas growth on 25% PEG resulted in reduced transpiration rate in non-colonized plants, SA190-colonized plants showed a much higher transpiration rate, consistent with a better overall water status under stress conditions (Fig. 5G).

Stomatal closure is a common adaptation response of plants to drought conditions to reduce water loss. We therefore analyzed stomatal opening under non-stress and 25% PEG in SA190-colonized and non-colonized plants (Fig. 5H). Under non-stress growth conditions, no statistically significant differences in stomatal aperture were observed between SA190-colonized and non-colonized plants (Fig. EV5B). However, under drought stress conditions, non-colonized plants rapidly induced stomatal closure, whereas SA190-colonized plants showed significantly more open stomata (Fig. 5I).

The observation that SA190-colonized plants showed higher water contents and significantly higher growth rates under water limiting conditions suggested that SA190 plants might make better use of the available water. We therefore measured whether SA190 altered the water use efficiency (WUE) of its host plant. We observed no significant change in water use efficiency between mock- and SA190-colonized plants under normal conditions (Fig. EV5C). However, when plants were exposed to drought stress, the water use efficiency of SA190-colonized plants was clearly superior to that of non-colonized plants (Fig. 5J). Overall, these results suggest that an enhanced water status in SA190-colonized plants is the basis for maintaining growth of Arabidopsis plants under limiting water conditions.

## Discussion

Water is critical for life on earth and is a major determinant of food production worldwide. Due to the increasing world population and climate change, water is becoming increasingly scarce on our planet. Since globally 70% of all freshwater is used by agriculture, increasing the water use efficiency of crops as well as making crops more drought tolerant are among the most important targets in agricultural engineering. Since water uptake primarily occurs via roots, strategies to improve water use efficiency through changes in root system architecture have recently gained an increased interest. Besides genetic engineering and molecular breeding, the application of PGPBs might offer another way to modify the root system and enhance water use efficiency and drought tolerance of crops. We report in this work that the root endophytic bacterium *Pseudomonas argentinensis* SA190 isolated from the roots of the desert plant *Indigofera argentea* induces major changes in root morphogenesis and maintains growth of plants under limiting water conditions.

Tolerance to environmental stresses can be achieved via different modes of cellular and biochemical changes conferred by beneficial microbes. Among the physiological and molecular responses to drought conditions, the plant phytohormone ABA plants plays a central role in drought adaptation. ABA levels rapidly rise upon water limitation, resulting in stomatal closure to reduce transpiration and save water at the cost of growth (Zhu, 2002). Interestingly, under drought stress conditions, SA190-colonized plants showed an improved leaf water status and transpiration rate as well as maintaining plant growth. These experiments were conducted under maximally controlled conditions using the genetic model plant Arabidopsis. Since it frequently happens that crops behave completely different in truly agricultural conditions, we tested whether these findings can also be translated to an important agricultural production system of the fodder plant alfalfa in open field agriculture conditions. Field trials with SA190-inoculated alfalfa plants showed superior biomass production when compared to productivity of non-colonized plants. These results support the concept that beneficial microbes such as SA190 might be a powerful tool to sustain agriculture production during water limiting conditions and events.

The analysis of the interaction between SA190 and Arabidopsis in the context of limited water availability revealed many of the known players of drought stress tolerance in plants. In particular, we found many genes involved in the ABA pathway and our subsequent genetic analysis showed that ABA is essential for mediating SA190-induced drought tolerance. Transcriptome and qRT-PCR analysis showed that at least seven aquaporin genes are differentially regulated in SA190-colonized plants under drought stress and this regulation by SA190 was found to be entirely dependent on the ABA pathway. Because aquaporins play a key role in adjusting water relations in plants and many are regulated by ABA (Jang et al, 2004), we used these genes as a proxy for the set of SA190-regulated ABA-dependent genes. Intriguingly, expression of all identified aquaporin genes was strongly suppressed under drought stress conditions in non-colonized plants, but none of these aquaporin genes were found to have significantly altered expression under non-stress conditions in SA190-colonized or non-colonized plants. These results suggested a potentially epigenetic priming mechanism. Since H3K4me3 enrichment is associated with transcriptionally active gene regions of drought responsive genes (van Dijk et al, 2010), we investigated the set of differentially regulated aquaporin genes in the context of SA190 colonization. We found that SA190 primes aquaporin genes by enhancing H3K4me3 levels in the regions near the transcriptional start sites. Priming by SA190 occurs already in the absence of drought stress. Therefore, SA190 interaction functions via priming genes for enhancing their expression under water limiting conditions. The H3K4me3 mark is broadly distributed on many ABA inducible genes (van Dijk et al., 2010) and mutants in the key enzyme of the ABA biosynthesis pathway *NCED3* show enriched H3K4me3 levels under drought stress conditions (Ding et al, 2011). This modification is mediated by the histone methyl transferase ATX1 (Arabidopsis trithorax-like 1), as transcript levels of several ABA and drought-upregulated genes like *RD29A* and *RD29B* were reduced during drought treatment in the *atx1* mutant (Ding et al, 2012b). It will be interesting to see if ATX1 and/or other methyl transferases are also involved in mediating H3K4me3 priming of SA190-targeted genes.

Aquaporins play a key role in transmembrane water transport in transpiration and environmental stress responses (Maurel et al., 2015). In Arabidopsis there exist 35 aquaporin genes (Johanson et al., 2001) which can be broadly divided into plasma membrane intrinsic proteins (PIPs) and tonoplast intrinsic protein (TIPs) genes. Some of the Arabidopsis PIPs and TIPs proved to be active water channels in *Xenopus* oocytes (Quigley et al., 2002).The biological significance of aquaporins in plants is their ability to modulate transmembrane water transport in situations where adjustment of water flow is physiologically critical (Li et al., 2014). PIPs play an important role in controlling the transcellular water transport and are subdivided into the two subfamilies PIP1 and PIP2 (Maurel et al., 2015). The overexpression of PIP isoforms increased the root osmotic hydraulic conductivity, transpiration and shoot to root ratio while the downregulation leads to drought stress susceptibility (Pawłowicz & Masajada, 2019). However, when expressed in heterologous systems, overexpression of aquaporins can also have negative effects on stress resistance, due to the fact that the native stress response machinery may recognize them as a foreign proteins (Li et al., 2014). Aquaporins are also regulated by post-translational modification including phosphorylation, methylation, acetylation, glycosylation, and deamination. Although fine-tuning aquaporin expression plays a key role in all functions related to the water status of plants, the multitude of genes and their potential redundancy makes genetic analyses a complicated matter. Redundancy of aquaporins in Arabidopsis might also be due to limiting our success in analyzing aquaporin mutants with respect to SA190-induced drought stress tolerance (Fig. 5B). However, using aquaporin inhibitors (AgNO_3_ and HgCl_2_), although probably not highly specific, independently suggested that aquaporins might mediate part of SA190-induced drought stress tolerance (Fig. 5D). These results require further research to unequivocally clarify the role of aquaporins in SA190-induced drought tolerance.

Drought stress massively alters many physiological parameters in plants, with ABA playing one of the primary roles in abiotic stress tolerance (Nakashima & Yamaguchi-Shinozaki, 2013; Zhu, 2002). A cluster of ABA-related genes was found in the transcriptome of SA190-colonized plants. Our genetic analysis showed that the ABA pathway is essential for mediating SA190-induced drought tolerance by priming of target genes. We conclude that SA190-induced drought tolerance is mediated by enhancing H3K4me3 levels target genes in an ABA-dependent manner (Fig. EV6). Priming by beneficial microbes bears an enormous potential in agricultural engineering which suffers from the problem that expressing stress-related genes often comes with a cost in growth and yield. Given that desert plant-associated beneficial microbes, such as *Enterobacter sp*. SA187 can enhance salt and heat tolerance in plants (de Zelicourt et al., 2018; Shekhawat et al, 2021), desert beneficial microbes, such as SA190, might present an affordable and readily available technology to ensure food production in arid countries with limited water availability. On the other hand, considering the effects of global climate change on, desert beneficial microbes might also help to limit yield losses in less arid regions suffering from intermittent drought and heat stress periods.

## Materials and Methods

### Plant material, seedling colonization, and stress assays

Seeds of *Arabidopsis thaliana* ecotype Col-0 were surface sterilized for 10 min in 70% ethanol with 0.05% Triton X-100, washed three times with 100% ethanol and dried in a laminar flow hood. Sterilized seeds were scattered on half strength MS (½MS) medium supplemented with either 100 μl of Luria Broth (LB) medium (Mock) or a fresh culture of SA190 grown in LB to an optical density of 0.21, at final concentration of 10^8^ cfu/ml. Seeds were stratified in the dark for 2 days at 4°C and then transferred in a growth chamber set to 22 °C with a long-day photoperiod (16h light, 8h dark) for 5 days. The germinated seedlings (~1.0–1.5 cm root length) were then transferred to ½MS as a normal condition or to ½MS infiltrated with 25% Polyethylene-glycol (PEG) 8000 (Fisher Scientific, Belgium) to induce drought stress. Six seedlings were used per square petri plate. The number of lateral roots (LR) was evaluated under a stereomicroscope and the root length was measured using ImageJ on the 9th day of stress treatment. Lateral root density (LRD) was calculated by dividing the number of lateral roots by the primary root length. The fresh weight (FW) of shoots and roots were taken after 16 days of stress. Dry weight (DW) was measured after drying the shoot and root tissues for 2 days at 80°C. All assays were performed in three biological replicates (bacterial colonies) and two technical replicates (petri plates). The following mutant lines were used in this study: abscisic acid mutants (*aba2-1, pyr1pyl4,5,8*), ethylene mutant (*ein2-1, ein3-1 and acs1-1*) and Aquaporin mutants *tip2;1* and *pip1;1 pip1;2 pip1;3 pip1,4 pip1;5* (q*pip*). Aquaporin inhibitors AgNO_3_ (silver nitrate, Sigma) and HgCl_2_ (mercury chloride, Sigma) were added to pre-cooled ½MS agar medium together with 25% PEG.

### Laser scanning confocal microscopy imaging

The SA190:GFP strain was generated as described in (de Zelicourt et al., 2018). Briefly, the GFP was introduced by the *E.coli* SM10λpir strain carrying the GFP donor plasmid, pUX-BF13 and the pRK600 mobiliser plasmid. To determine the colonization of SA190:GFP on plant tissue the 5 day old seedlings, mounted in 100 μg/ml Propidium Iodide (PI), were used and visualized using confocal microscope ZEISS LSM880 with Airyscan with the Plan-Apochromat 10× (n.a. 0.45) objective lense for small magnifications (100 nm) or the Plan-Apochromat 63x (n.a 1.4) oil immersive objective lense for higher magnifications (20 nm) For excitation of both GFP and PI, we used the argon-based laser with power output 6.5 or 9 % for 10× or 63× objective lense, respectively and samples were excited at wavelength 493-584 nm for GFP and 604-718 nm for PI. The ZEN black edition 3.0 SR was used for assembly of images and the ZEN blue lite edition 3.0 was used for image editing.

### Bacterial colonization

Shoots and roots of 21-days old mock- and SA190-colonized shoots of Arabidopsis grown on ½ MS with or without 25% PEG were collected in Eppendorf tubes. Fresh weight was recorded using sensitive balance (METTLER TOLEDO). Samples were grinded using Qiagen Tissue Lyser II Sample (Disruption-11843) for 2 minutes with 500 μl of extraction buffer (10mM MgCl2+ 0.01% silwet 77), and then were incubated for 1h at 28°C with shaking at 300 rpm (Eppendorf, ThermoMixer C). Samples were diluted 10-fold, and then spread on LB agar and colony forming units (CFUs) were counted after overnight incubation at 28°C. Calculated number of CFUs was normalized to plant fresh weight.

### Stomatal aperture measurement assay

To measure the stomatal parameters, 5d old seedlings inoculated with +/-SA190 on ½MS were transferred and grown further for 16 days on ½MS +/-25% PEG plates. The leaves from same developmental stage were excised from the plants and a section from middle part of the lamina excluding the mid rib was cut and mounted on a double sided tape, sticked to a microscopic glass slide, with the abaxial surface in contact with the tape. The upper green tissues were then scraped off by scalpel and the remaining lower epidermis was imaged using an Axio Imager Z2 microscope (Zeiss) equipped with DIC optics and EC Plan-Neofluar. For measuring the stomatal aperture, the aperture width and total stomatal length of each stoma were manually measured using ImageJ and their ratios were analyzed statistically. A minimum of 30 stomata was measured from each leaf. The experiment was repeated thrice and samples were prepared from at least 3 leaves from each genotype per biological replicate.

### Transpiration changes

To conduct the transpiration rates assays, the rosettes of 21-days old mock- and SA190-colonized shoots of Arabidopsis grown on ½ MS with or without 25% PEG were weighed at regular 5 min intervals. The transpiration rate was calculated using the formula: WL (mg) = FWi (initial FW) − desiccated weight (Cohen et al, 2015).

### Leaf relative water content

The 4^th^ leaf from the rosette of six 21-days old non-colonized or SA190-colonized plants treated with 0 or 25% PEG was used to measure the relative water content (RWC). After measuring the fresh weight, leaves were placed in distilled water for 3 hours in the dark and the turgid weight was recorded. The samples were oven-dried at 70°C for 24h and the dry weight was measured according to the formula: LRWC (%) = [(FW-DW)/(TW-DW)]×100 (Sade et al, 2015).

### Water Use Efficiency (WUE)

To determine the WUE for plants, sterilized 50 mL tubes were filled with a sterilize soil-perlite mixture (w:w 1:1) and 35 ml of water was added to each tube. Single 5 day old Arabidopsis seedling that had been cultured on plates with or without SA190 were transferred to the hole in the lid of individual tubes. The plants were treated as described in (Wituszynska & Karpiński, 2014). For the drought condition samples, the plants were cultivated for an extra 14 days.

### Drought assays in soil

5-days old mock- and SA190-colonized Arabidopsis seedlings were transferred to jiffy pots in four biological replicates of 9 plants each. The plants were grown in Percival growth chambers and watered twice a week for two weeks, then watering was stopped for 3 weeks to induce drought conditions, before rewatering again for 7 days to allow the plants to recover. The plants were photographed at each step using Canon EOS 6D.

### RNA extraction

Total RNA was extracted from the roots of 21-days old mock- and SA190-colonized Arabidopsis grown on ½ MS with or without 25% PEG with the Nucleospin RNA plant kit (Macherey-Nagel) following the manufacturer’s recommendations. The quality and quantity of the RNA was assessed using Nanodrop-6000 spectrophotometer, 2100-Bioanalyzer (RNA integrity number greater than 8.0) and QubitTM 2.0 Fluorometer with the RNA BR assay kit (Invitrogen).

### RNAseq Library Construction and Sequencing

In order to prepare the cDNA library, we used 1 μg total RNA per sample. The ribosomal RNA was removed using a Ribo-Zero Magnetic Kit with a 1:1 mixture of Ribo-Zero Magnetic Kit (Bacteria) and Ribo-Zero Magnetic Kit (Plant) following the manufacturer’s recommendations. The quality of the library was assessed on the Agilent Bioanalyzer 2100 system. Sequencing was performed using Illumina HiSeq deep sequencing (Illumina HiSeq 2000, Illumina).

### Transcriptome analysis

We performed transcriptome sequencing for each library of Arabidopsis to generate 101-bp paired-end reads on Illumina HiSeq4000 Genome Analyzer platform. Low quality reads were trimmed using the Trimmomatic version 0.32 (Bolger et al, 2014) (http://www.usadellab.org/cms/?page=trimmomatic) with the following parameters: Minimum length of 36 bp; Mean Phred quality score greater than 30; Leading and trailing bases removal with base quality below 3; Sliding window of 4:15. After pre-processing the Illumina reads, the reads are mapped on to transcripts using TopHat (Trapnell et al, 2009) (ver. 2.1.1; http://tophat.cbcb.umd.edu/) for aligning with the genome. For TopHat, the Reference-Arabidopsis thaliana (TAIR10) genome (https://www.arabidopsis.org) was used as the reference sequences with maximum number of mismatches as 2. We used count-based normalization implemented in DESEQ2 (Love et al, 2014). To quantify the reads from the mapped alignment, featureCounts package was overlapped with each gene co-ordinates (Liao et al, 2013).

To identify the differentially expressed genes, the following parameters were used: p-value of 0.05 with a statistical correction using Benjamini Hochberg FDR of 0.05 in DESEQ2. A cut-off of 2 fold up- or down-regulation has been chosen to define differential expression. After processing the data, visualization of differential expression was done using cummerbund v2.14.0 (http://bioconductor.org/packages/release/bioc/html/cummeRbund.html).

Hierarchical clustering of the quantified genes were performed using MeV 4.9.0 version (TM4, https://sourceforge.net/projects/mev-tm4/files/mev-tm4/MeV%204.9.0/) utilizing the Pearson correlation method. Enriched GO term analysis was carried out using DAVID (Huang et al, 2009; Sherman et al, 2022).

### qRT-PCR analysis

Total RNA was extracted from 21-days old mock- and SA190-colonized Arabidopsis grown on ½ MS +/-25% PEG using NucleoSpin Plant RNA (Macherey Nagel) kit following the manufacturer’s protocol. First strand cDNA was synthesized from 1μg of total RNA using SuperScript III First-Strand Synthesis SuperMix kit (Invitrogen). The diluted cDNA was used to perform quantitative RT-PCR (qRT-PCR) using SsoAdvanced Universal SYBR Green Supermix (Bio-Rad). All reactions were amplified in a CFX96 Touch Real-Time PCR Detection System (BIO-RAD) at 50°C for 2 min, 95°C for 10 min, and 40 cycles of 95°C for 10 sec and 60°C for 40 sec, followed by a dissociation step to validate the PCR products. The data was analyzed using Bio-Rad CFX manager software. Tubulin was used as a housekeeping gene for normalization of gene expression levels. Analyses were performed in triplicate and were repeated thrice with independent RNA samples. Primers used in this study are listed in Supplementary table 1.

### Chromatin Immunoprecipitation (ChIP) qPCRs

We conducted ChIP as described in previous studies (Shekhawat et al., 2021). Briefly, roughly 1g of 21-days old mock- and SA190-colonized Arabidopsis grown on ½ MS with or without 25% PEG were cross-linked by vacuum-infiltrating 1% formaldehyde for 15 min and subsequent quenching by 2 M glycine. Nuclei were extracted from the frozen ground powder using NIB (0.4 M Sucrose, 10 mM Tris-HCl pH8, 10 mM MgCl2, 5 mM ß-mercaptoethaol and 1x Proteases Inhibitor cocktail). Nuclei were lysed in NLB (50 mM Tris-HCl pH8, 10 mM EDTA, 1% SDS and 1x Proteases Inhibitor cocktail. Chromatin was sonicated using a Diagenode Bioruptor (40 Hz, 14 cycles each with 30s on/30 s off with ice cooling), yielding fragments with a size of around 250-350 bp. Antibodies (anti-H3, ab1791; anti-H3K4me3, from Abcam, ab8580 http://www.abcam.com) were incubated with protein A-coated agarose beads (Invitrogen) for at least 2 hour at 4°C in IP buffer (1.1 % Triton X-10, 1.2mM EDTA, 16.7 mM Tris-HCl pH8, 167 mM NaCl) and 1x Proteases Inhibitor cocktail. Immunoprecipitations were done in IP buffer at 4°C for overnight. After washing with low (150 mM NaCl) and high salt (500 mM NaCl) IP buffers and reverse crosslinking, resulted DNA was extracted using the phenol–chloroform method and precipitated with ice chilled ethanol and glycogen (Invitrogen), and then re-suspended in 20 μl of water. ChIP-PCR was performed for three regions of indicated gene loci. Amplification values were normalized to H3 (normalized signal modification/normalized signal H3). The given values in graphs are the means of two biological replicates, with each replicate was normalized to the respective Col-0 with no treatment (mock) sample before averaging.

### Field trials

Open field trials were conducted at the experimental station in Hada Al-Sham (N 21°47’47.1”, E 39°43’48.8”) Saudi Arabia, in the winter seasons 2015–2016. The experiments were performed in a randomized complete block design with a split-split plot arrangement of four replicates, plots (2 × 1.5 m) with seed spacing 20 cm row-to-row. The field was irrigated using groundwater as full irrigation or by a 25% reduction of irrigation. The soil had an average pH 7.74 and salinity EC = 1.95 dS·m-1. The yield was recorded every 25-30 days from each harvest; three harvests were done. Field trials data were analyzed as a randomized complete block design, the analysis of variance (one-way ANOVA) of four replicates, was performed and LSD (Duncan’s test, P< 0.05) was calculated to test the significance of differences between means. To inoculate alfalfa (Medicago sativa var. CUF 101) seeds was done according to (Daur et al, 2018). In brief, a slurry was prepared consisting of sterilized peat, a broth culture of SA190, and sterilized sugar solution (10%) in the ratio 5:4:1 (w:v:v). Subsequently, alfalfa seeds were coated with the slurry at a rate of 50 mL/kg. As a control, seeds were coated with a similar mixture without bacteria.

## Data Availability

The raw data of RNA-sequencing have been deposited in NCBI with GSE Number GSE184355. Any additional information required to reanalyze the data reported in this study is available from the lead contact upon request.

## Acknowledgements

The work was supported by a grant to HH from the King Abdullah University of Science and Technology BAS/1/1062-01-01. We thank Pedro Rodriguez for providing *aba2-1* and *qpylpyr* mutants. We thank KAUST core lab facility for RNA seq and hormone measurements processes. We thank Dr. Sabiha Parveen for helping with data uploading. We would like to thank all members of the Hirt lab for useful discussion.

## Author Contributions

HH conceived and directed the research. KMA, AAR and AHS performed most of the lab experiments. ID, MMS performed open field experiments. AV performed RNA seq analysis. MK, RJ, KF, NT, WA helped with many experiments. MMS and AHS coordinated the research. HH, AHS, AAR, KMA, and MMS wrote the manuscript with inputs from other authors.

## Conflict of Interest

The authors declare no conflict of interest.

## Expanded View Figures

**EV1.**
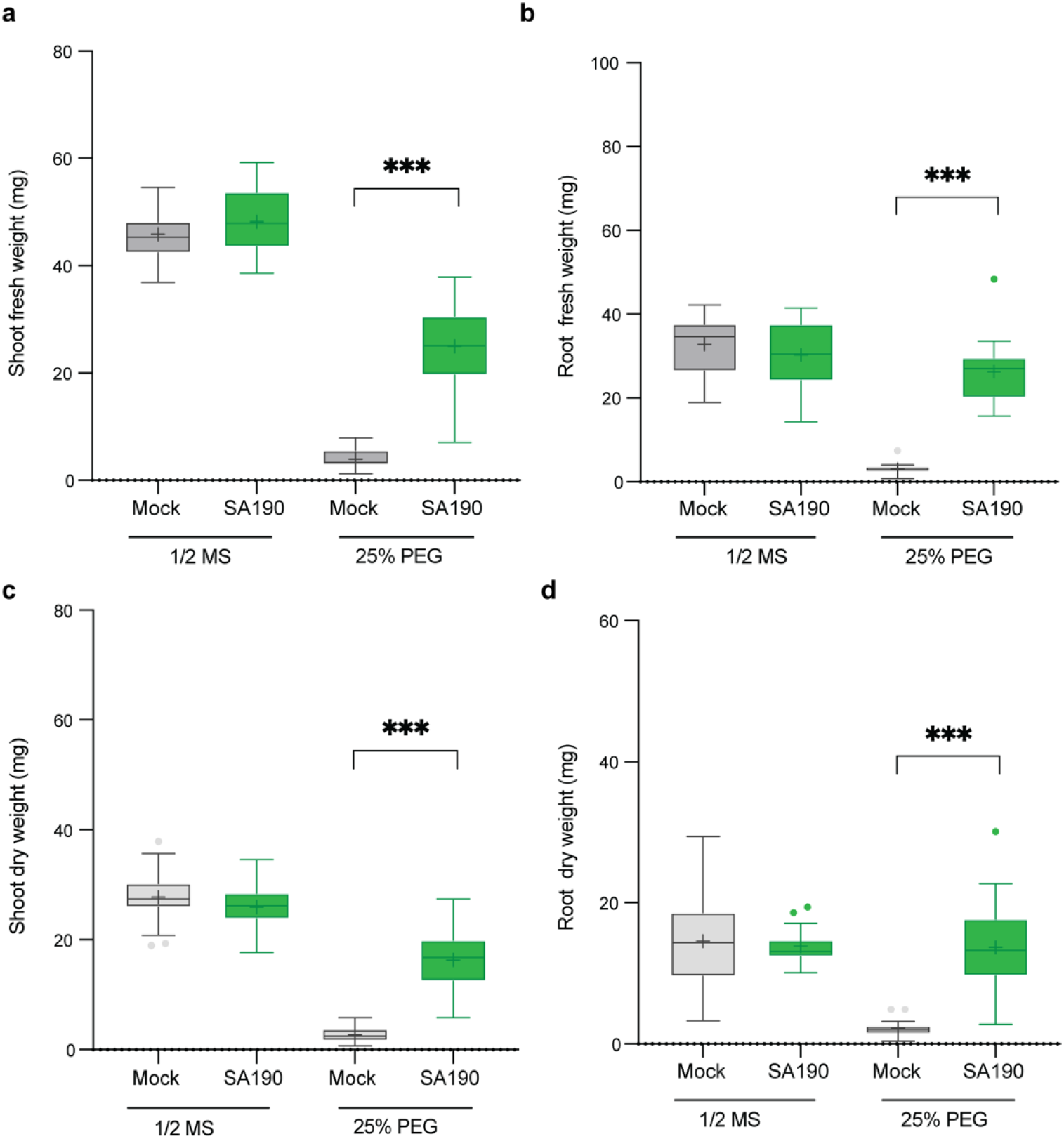
SA190 enhances Arabidopsis drought stress tolerance. **A.** Shoot fresh weight and **B.** Root fresh weight (b) of 16-day-old mock- and SA190-colonized plants grown on ½ MS medium and ½ MS+PEG medium. **C.** Shoot dry weight and **D.** Root dry weight of 16-day-old mock- and SA190-colonized plants grown on ½ MS medium and ½ MS+PEG medium. All plots represent the mean of 3 biological replicates (n > 39). Error bars represent SE. Asterisks indicate the statistical differences based on the Student’s t-test (* P < 0.05; ** P < 0.01; *** P < 0.001).

**EV2.**
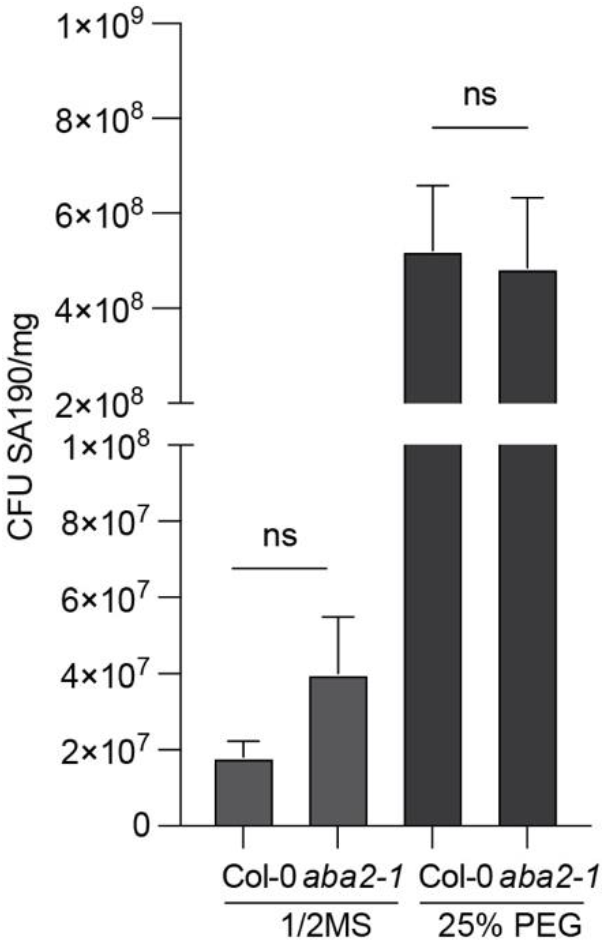
Efficiency of root colonization by SA190 in 21 days old Col-0 and *aba2-1* plants grown for 16 days under normal (½ MS) or drought conditions (½ MS + 25% PEG).

**EV3.**
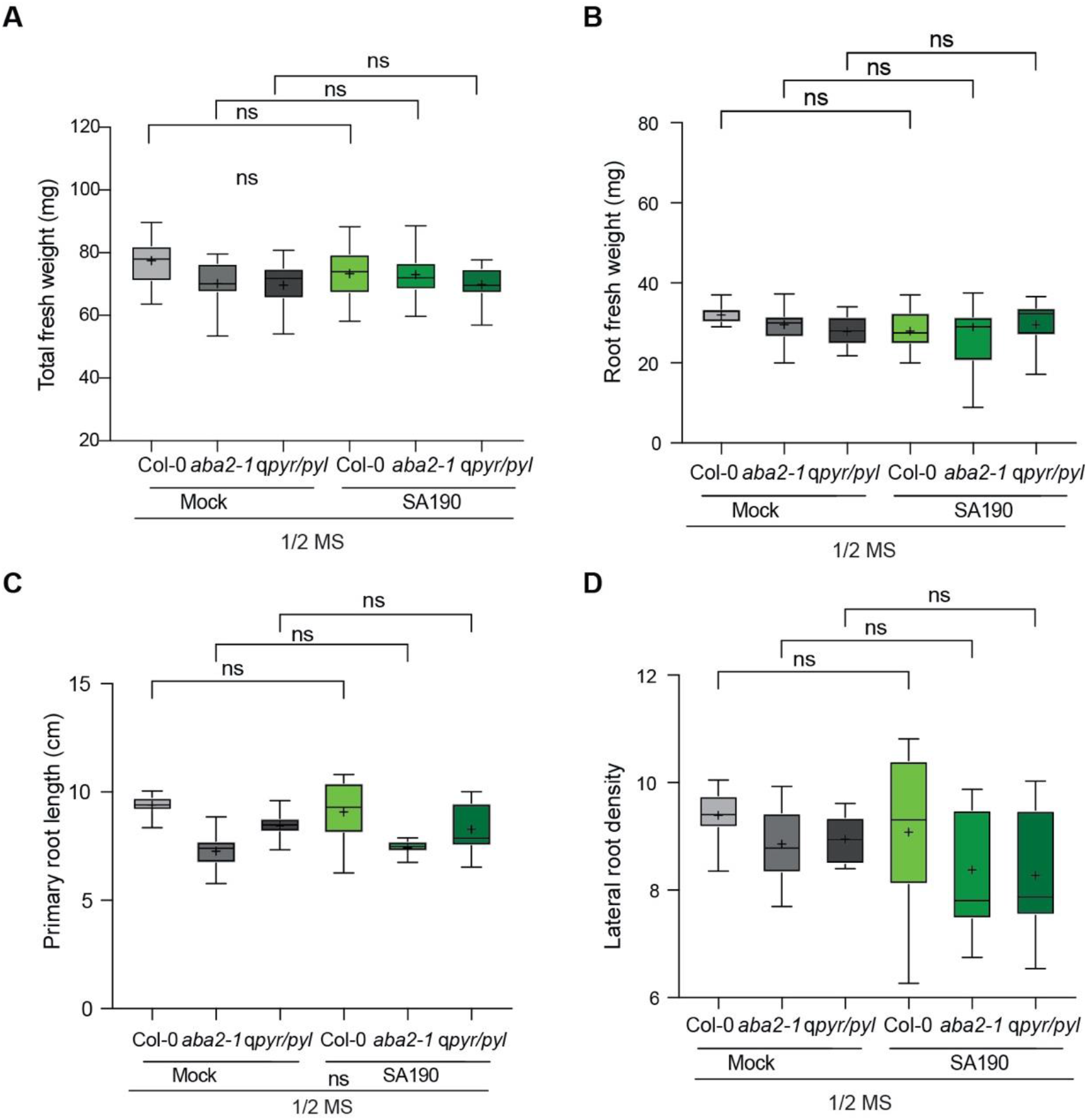
**A-B.** Total fresh weight and root fresh weight of 21 days old non-colonized- (Mock) and SA190-colonized (SA190) Col-0, *aba2-1* and *qpyr/pyl* plants grown under non stress conditions (½ MS). **C-D.** Primary root length and Lateral root density of 21-day-old, mock- and SA190-colonized Col-0, *aba2-1* and *qpyr/pyl* plants upon growth for 16 days on ½ MS plates

**EV4.**
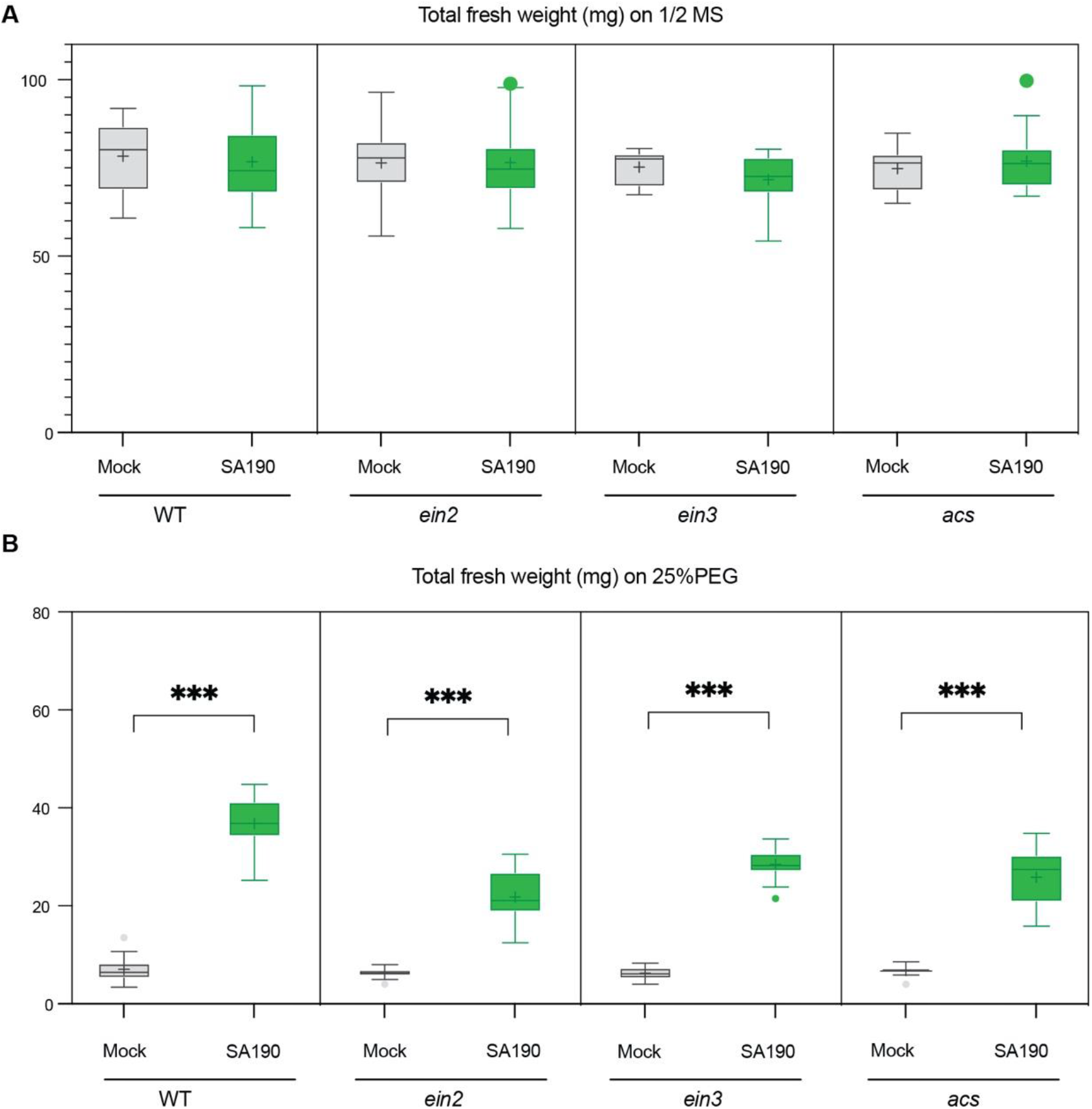
Arabidopsis-SA190 interaction is independent of the ethylene pathway. Total fresh weight of mock- and SA190-colonized 16-day-old ethylene mutants on **(A)** ½ MS or **(B)** 25%PEG. Error bars represent SE and asterisks indicate a statistical difference based on the Student’s t-test (* P < 0.05; ** P < 0.01; *** P < 0.001).

**EV5.**
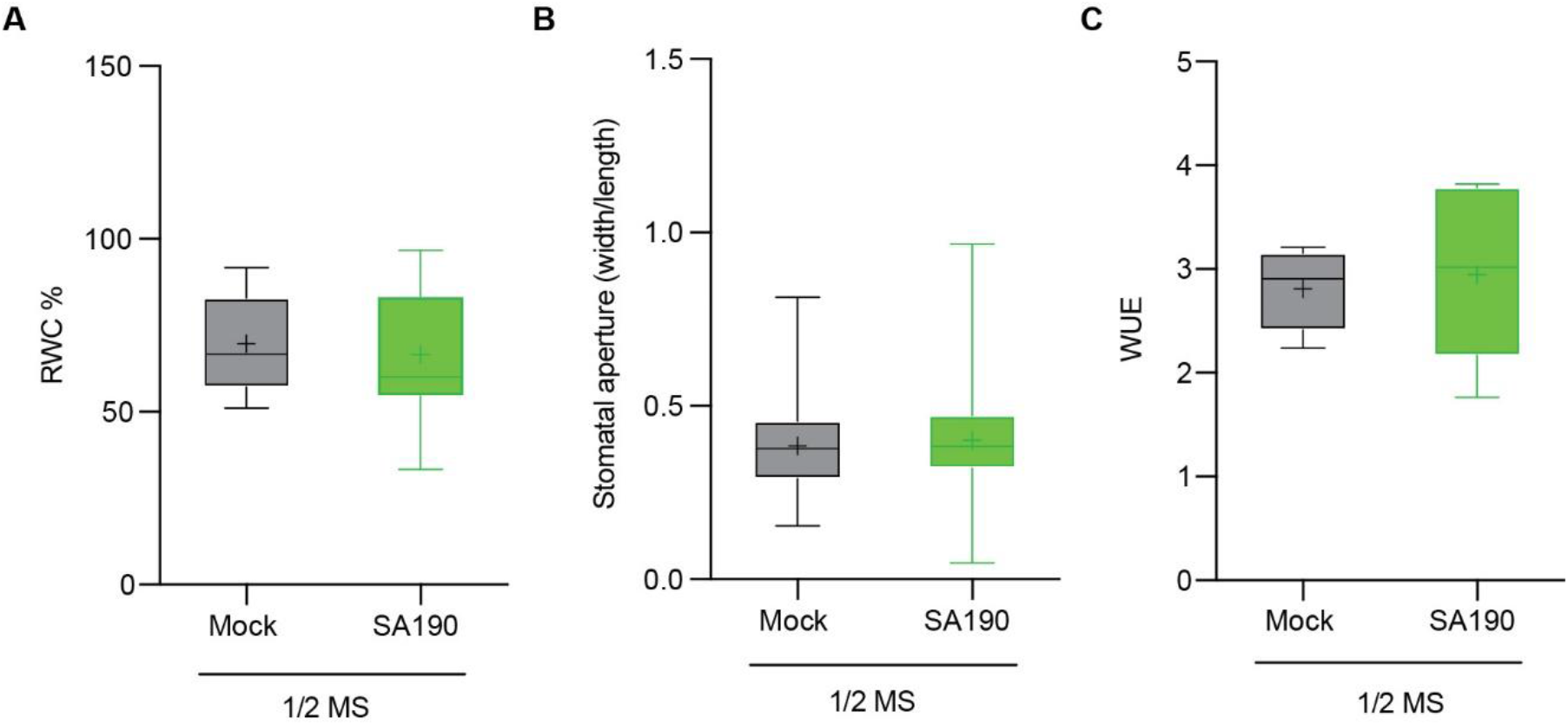
Role of SA190 colonization in plant drought stress physiology. **A.** The percentage of leaf relative water content (RWC) of 21-days old mock- or SA190-colonized plants grown for 16 days under normal (½ MS) conditions. The RWC was calculated as the formula RWC (%) = [(FW-DW)/(TW-DW)]x100, where FW is fresh weight, DW is dry weight and TW is the weight of fully turgid leaves. **B.** Comparison of stomatal aperture of SA190- and non-colonized 21-days old plants grown for 16 days under non-stress (½ MS) conditions. Scale bar = 20 μm. **C.** Water use efficiency (WUE) of 4-weeks old mock- or SA190-colonized plants grown under non-stress (½ MS) conditions. WUE was calculated as mg of dry weight produced per ml of water used. All plots represent the mean of 3 biological replicates. Error bars represent SE. Asterisks indicate a statistical difference based on the Student’s t-test (* P < 0.05; ** P < 0.01; *** P < 0.001).

**EV6.**
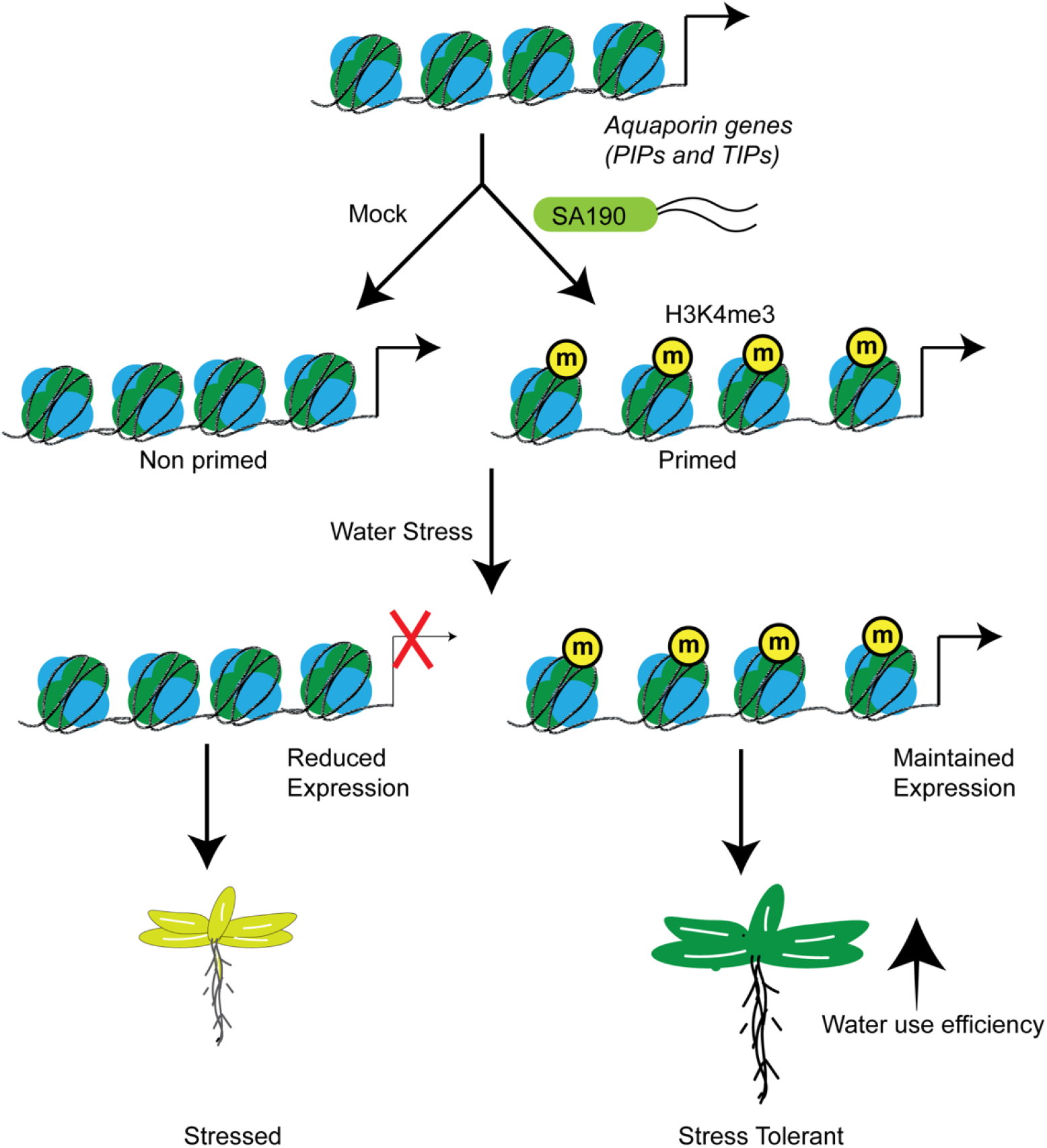
Graphical model of SA190 mediated aquaporin priming. SA190 constitutively deposits H3K4me3 histone marks on aquaporin promoters in an ABA-dependent manner to mediate gene expression for enhancing water use efficiency and drought stress tolerance.

---

**Supplementary Table 1.**
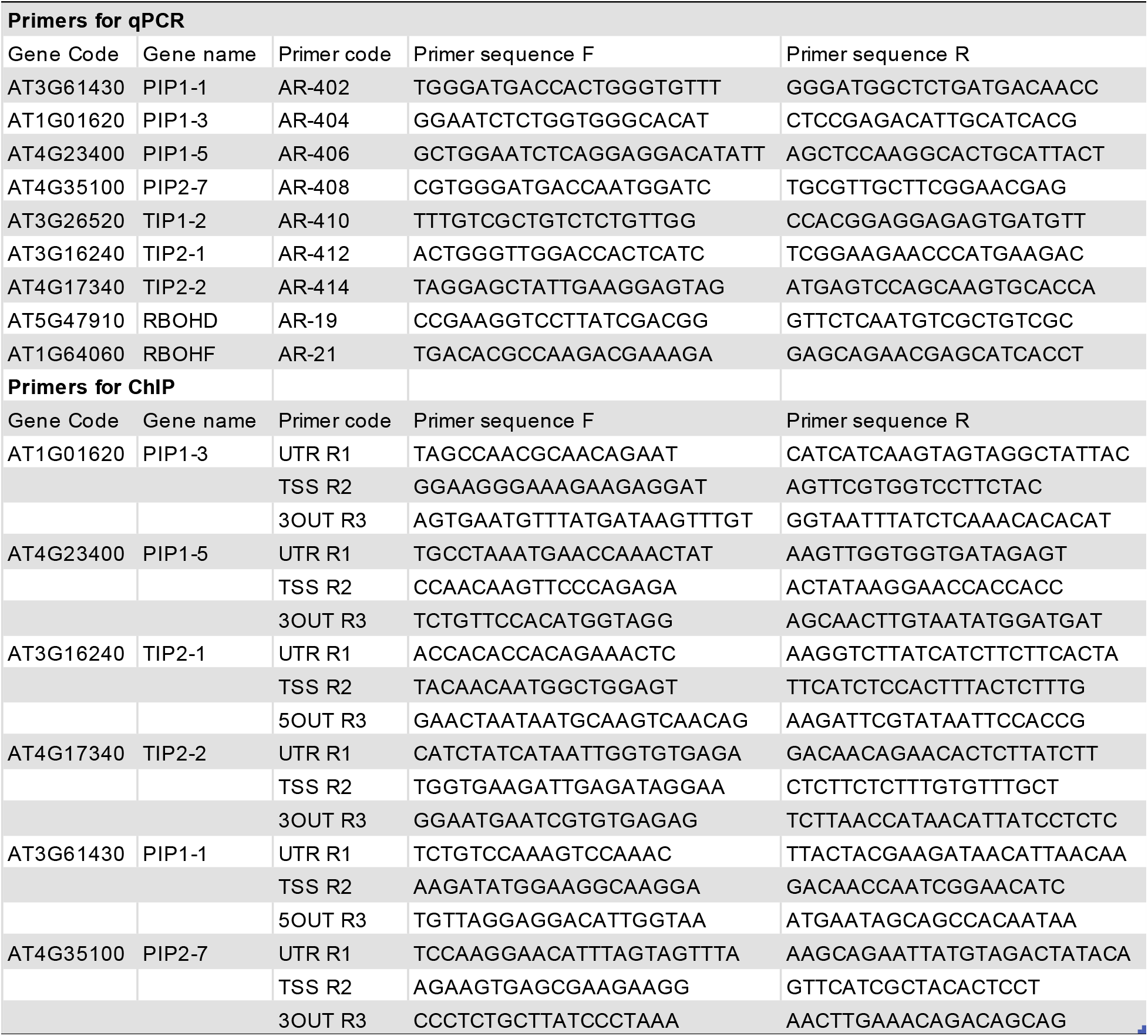

